# A comprehensive genetic characterisation of the mitochondrial Ca^2+^ uniporter in *Drosophila*

**DOI:** 10.1101/458174

**Authors:** Roberta Tufi, Thomas P. Gleeson, Sophia von Stockum, Victoria L. Hewitt, Juliette J. Lee, Ana Terriente-Felix, Alvaro Sanchez-Martinez, Elena Ziviani, Alexander J. Whitworth

## Abstract

Mitochondrial Ca^2+^ uptake is an important mediator of metabolism and cell death. Identification of components of the highly conserved mitochondrial Ca^2+^ uniporter has opened it up to genetic analysis in model organisms. Here we report a comprehensive genetic characterisation of the known uniporter components conserved in *Drosophila*. While loss of *MCU* or *EMRE* abolishes fast mitochondrial Ca^2+^ uptake, this results in surprisingly mild phenotypes. In contrast, loss of the regulatory gatekeeper component *MICU1* has a much more severe phenotype, being developmental lethal, consistent with unregulated Ca^2+^ uptake. Mutants for *MICU3* are viable with mild neurological phenotypes. Genetic interaction studies reveal that *MICU1* and *MICU3* are not functionally interchangeable. More surprisingly, loss of *MCU* or *EMRE* does not suppress *MICU1* mutant lethality, suggesting that the lethality results from MCU-independent functions. This study helps shed light on the physiological requirements of the mitochondrial Ca^2+^ uniporter, and provides a suite of tools to interrogate their interplay in homeostasis and disease conditions.

## Introduction

The uptake of Ca^2+^ into mitochondria has long been established as a key regulator of an array of cellular homeostatic processes as diverse as bioenergetics and cell death [1,2]. A series of seminal discoveries have recently elucidated the identity of the components that make up the mitochondrial Ca^2+^ uniporter complex. The mammalian uniporter is comprised of MCU as the main pore-forming protein [3,4], its paralog MCUb [5], a small structural component, EMRE [6], and the regulatory subunits, MICU1-3 [7-9]. Reconstitution studies in yeast, which lacks a mitochondrial Ca^2+^ uniporter, have demonstrated that heterologous co-expression of MCU and EMRE are necessary and sufficient to confer uniporter activity [10]. The family of EF-hand-containing proteins, MICU1, MICU2 and MICU3, have been shown to exhibit a ‘gatekeeper’ function for the uniporter, serving to inhibit Ca^2+^ uptake at low cytoplasmic concentrations [9,11,12]. These components are generally highly conserved across eukaryotes, including most metazoans and plants, but not in many fungi and protozoans, reflecting their ancient and fundamental role [13].

While the composition and function of the uniporter have been well-characterised *in vitro* and in cell culture models, the physiological role of the uniporter is beginning to emerge with *in vivo* characterisation of knockout mutants [14]. Current data present a complex picture. Initial studies of *MCU* knockout mice described a viable strain with a very modest phenotype, in a mixed genetic background [15], although subsequent studies using an inbred background reported *MCU* loss to be lethal or semi-viable [16] and tissue-specific conditional knockout revealed an important role in cardiac homeostasis [17]. Similarly, loss of *MICU1* in mice has a complex phenotype, varying from fully penetrant perinatal lethality [18] to incomplete lethality with a range of neuromuscular defects that unexpectedly improve over time in surviving animals [19].

One explanation for the reported phenotypic variability is that perturbing mitochondrial Ca^2+^ uptake can be influenced by additional factors, the most obvious being genetic background. Hence, there is a need for greater investigation into the physiological consequences of genetic manipulation of the uniporter components in a genetically powerful model system. Here we report a comprehensive genetic analysis of the uniporter complex components that are conserved in *Drosophila*. These include loss of function mutants for *MCU, EMRE, MICU1* and *MICU3* (*Drosophila* lack *MCUb* and *MICU2*), and corresponding inducible transgenic expression lines. Despite lacking fast Ca^2+^ uptake, *MCU* and *EMRE* mutants present a surprising lack of organismal phenotypes, although both mutants are short-lived, with a more pronounced effect when MCU is lost. In contrast, loss of *MICU1* causes developmental lethality, whereas mutants for *MICU3* are viable with modest phenotypes. Performing genetic interaction studies with these strains, we confirm the gatekeeper function of MICU1 is conserved in flies and reveal that *MICU1* and *MICU3* are not functionally interchangeable. More surprisingly, we find that loss of *MCU* or *EMRE* does not suppress *MICU1* mutant lethality, suggesting that the lethality results from MCU-independent functions. The generation of these genetic tools in *Drosophila* will facilitate the further investigation of the functional roles of the uniporter components *in vivo*.

## Results

To generate null mutants for *MCU* we used a P-element mobilisation technique exploiting a transposon at the 5’ end, MCU^EY01803^ (Fig. 1A). We isolated a single imprecise excision - a deletion of 1557 bp removing the *5’* end of *MCU* that includes the first three exons containing the start codon and mitochondrial targeting sequence common to most isoforms. We refer to this mutation as *MCU*^1^ (Fig. 1A). The neighbouring genes, *sulfateless* (*sfl*) and *javelin* (*jv*), remained intact and showed unaltered levels of expression (Fig. EV1A, B). The *MCU*^1^ deletion can be detected by genomic PCR and the breakpoints were verified by Sanger sequencing (Fig. 1B). Immunoblot analysis of crude mitochondrial extracts from homozygous *MCU*^1^ mutant homogenates using an antibody raised against the C-terminus of *Drosophila* MCU confirmed the absence of MCU protein (Fig. 1C).

**Figure 1.**
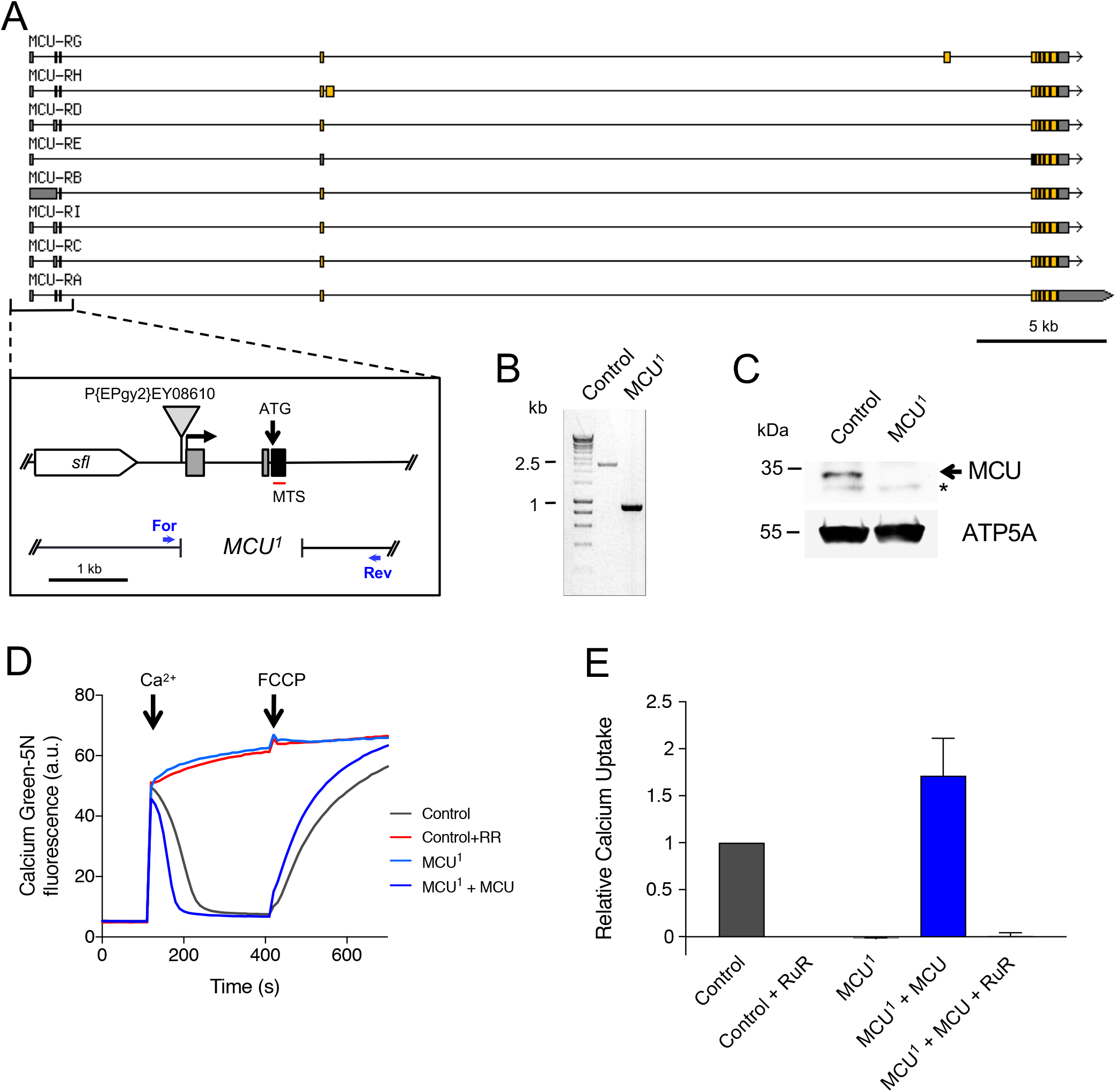
The *MCU^1^* Mutant Abolishes Fast Mitochondrial Ca^2+^ Uptake. **A.** Overview of *MCU* (CG18769) gene region (from FlyBase). The inset box details the 5’ region of *MCU*, including the neighbouring *sfl* (*sulfateless*). The P{EPgy2}EY08610 transposable element used to generate *MCU*^1^ is displayed, along with the location of the *MCU*^1^ breakpoints, and the forward and reverse screening primer positions. **B.** DNA analysis of *MCU*^1^. Primer sequences are detailed in the ‘Materials and Methods’ section. The control genotype yielded a ~2.5 kb band, compared to ~900 bp for *MCU*^1^ homozygotes. **C.** Western blot analysis of *MCU*^1^. Immunoblots were probed with the indicated antibodies. Asterisk denotes a non-specific band. Mitochondrial ATP5A is used as a loading control. **D.** Representative traces of Ca^2+^ uptake in mitochondria isolated from adult flies of the indicated genotypes after addition of 45 μM CaCl2. Extramitochondrial Ca^2^+ was measured by Calcium Green-5N fluorescence. Ca^2^+ was released from mitochondria by addition of 1 μM FCCP. *a.u.*: arbitrary units. The control genotype is *w*^1118^. Addition of the MCU inhibitor Ruthenium Red (RuR; 2 μM) blocks mitochondrial Ca^2^+ uptake, which is mirrored by *MCU*^1^. Mitochondrial Ca^2^+ uptake is restored by transgenic re-expression of *MCU*. **E.** Relative uptake kinetics were determined through linear fits of the initial phase of Ca^2^+ uptake and normalized to the wild type control (mean ± SEM, n = 3).

**Figure EV1.**
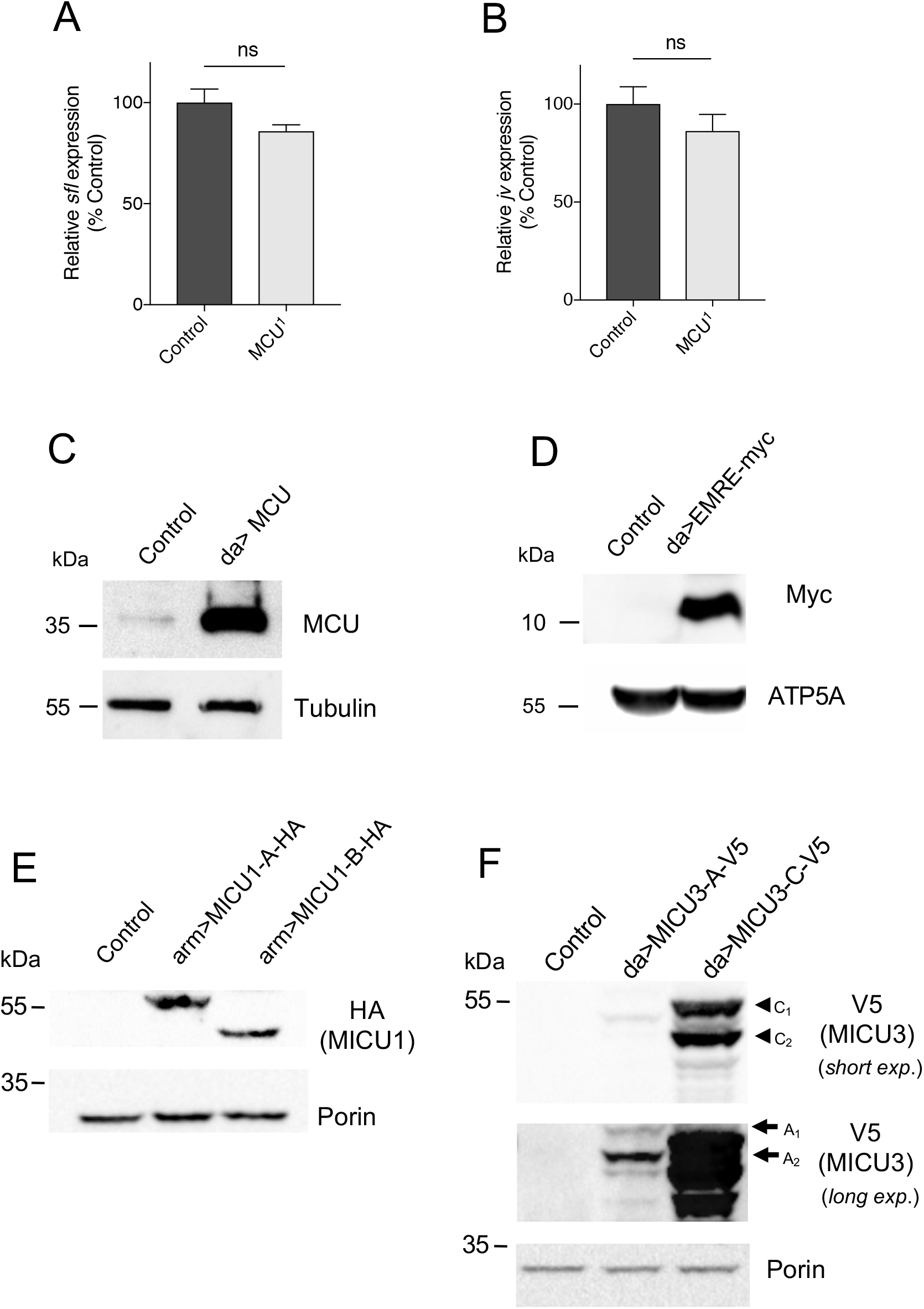
Expression analysis of *MCU* neighbouring genes and uniporter transgenes. **A, B.** Relative transcript level of *sfl* (**A**) or *jv* (**B**) for control and *MCU*^1^ flies (mean ± SD; n = 3). **C.** Western blot analysis of ubiquitously (*da-GAL4*) expressed *UAS-MCU* transgene. **D.** Western blot analysis of ubiquitously (*da-GAL4*) expressed *UAS-EMRE-myc* transgene. **E.** Western blot analysis of transgenic MICU1 expression of *UAS-MICU1-A-3xHA* and *UAS-MICU1-C-HA* lines using a ubiquitous (*arm-GAL4*) driver. **F.** Western blot analysis of relative MICU3 expression between UAS-MICU3-A-V5 and UAS-MICU3-C-V5 lines using a ubiquitous (*da-GAL4*) driver. In all conditions the control genotype was the relevant GAL4-only non-transgenic control (*GAL4/+*). Immunoblots were probed with the indicated antibodies.

Mitochondria from human or mouse cells lacking MCU fail to perform fast Ca^2+^ uptake [3,4,15]. To verify that *MCU*^1^ represents a functional null mutant, mitochondria were isolated from homozygous *MCU*^1^ adult flies and assayed for Ca^2+^ uptake. Similar to mammalian cells, the addition of Ca^2+^ to purified, energised mitochondria from wild-type *Drosophila* yields a rapid spike of extramitochondrial Calcium Green-5N fluorescence followed by a progressive decline in fluorescence as Ca^2+^ is buffered by mitochondria (Fig. 1D, E,). Ca^2+^ is released again upon depolarisation by the uncoupling agent FCCP, as reflected by the concomitant rise in Calcium Green-5N fluorescence. As expected, rapid Ca^2+^ uptake is completely blocked by the addition of the MCU inhibitor ruthenium red (RuR). This effect is fully replicated in *MCU*^1^ mutant mitochondria, reflecting a complete loss of fast Ca^2+^ uptake. Importantly, the lack of Ca^2+^ uptake was not due to loss of membrane potential as this was equivalent across the samples (EV2A, B). Moreover, Ca^2+^ uptake is restored upon transgenic expression of *MCU* (Fig. 1D, E, and EV2A, B). Together these data show that *MCU*^1^ is a null mutant incapable of fast Ca^2^+ uptake.

**Figure EV2.**
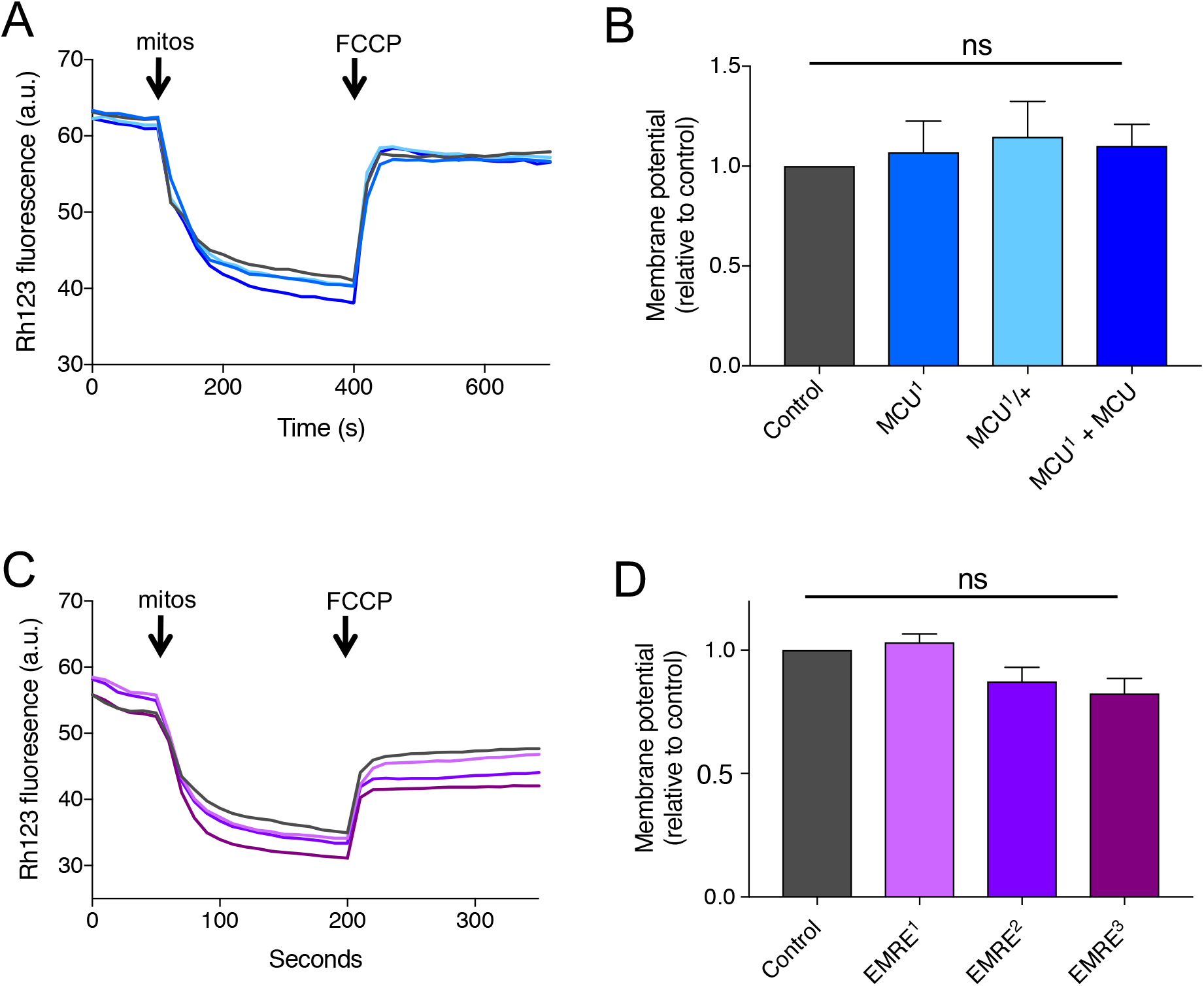
Analysis of mitochondrial membrane potential. **A, C.** Representative traces of Rhodamine 123 (Rh123) fluorescence (*a.u.:* arbitrary units) in isolated mitochondria of the indicated genotypes used for the Ca^2^+ uptake assays for (A) *MCU* (with *da-GAL4* expressed *UAS-MCU*) and (C) *EMRE* mutants (see Fig.1 and 3). **B, D.** Relative membrane potential was determined through Rh123 fluorescence change before and after addition of FCCP (ΔF-F0) normalized to controls (mean ± SEM; n = 3).

Despite this deficiency, *MCU*^1^ mutants are homozygous viable and develop to adult stage in expected Mendelian proportions (Fig. 2A). However, *MCU*^1^ mutants are significantly shorter-lived than controls (~34% reduction of median lifespan), a phenotype that is fully rescued by ubiquitous expression of transgenic *MCU* (Fig. 2B). Despite this attenuated longevity, *MCU*^1^ mutants do not display an appreciable defect in motor ability, as assessed via a negative geotaxis (climbing) assay, in young or older flies (Fig. 2C).

**Figure 2.**
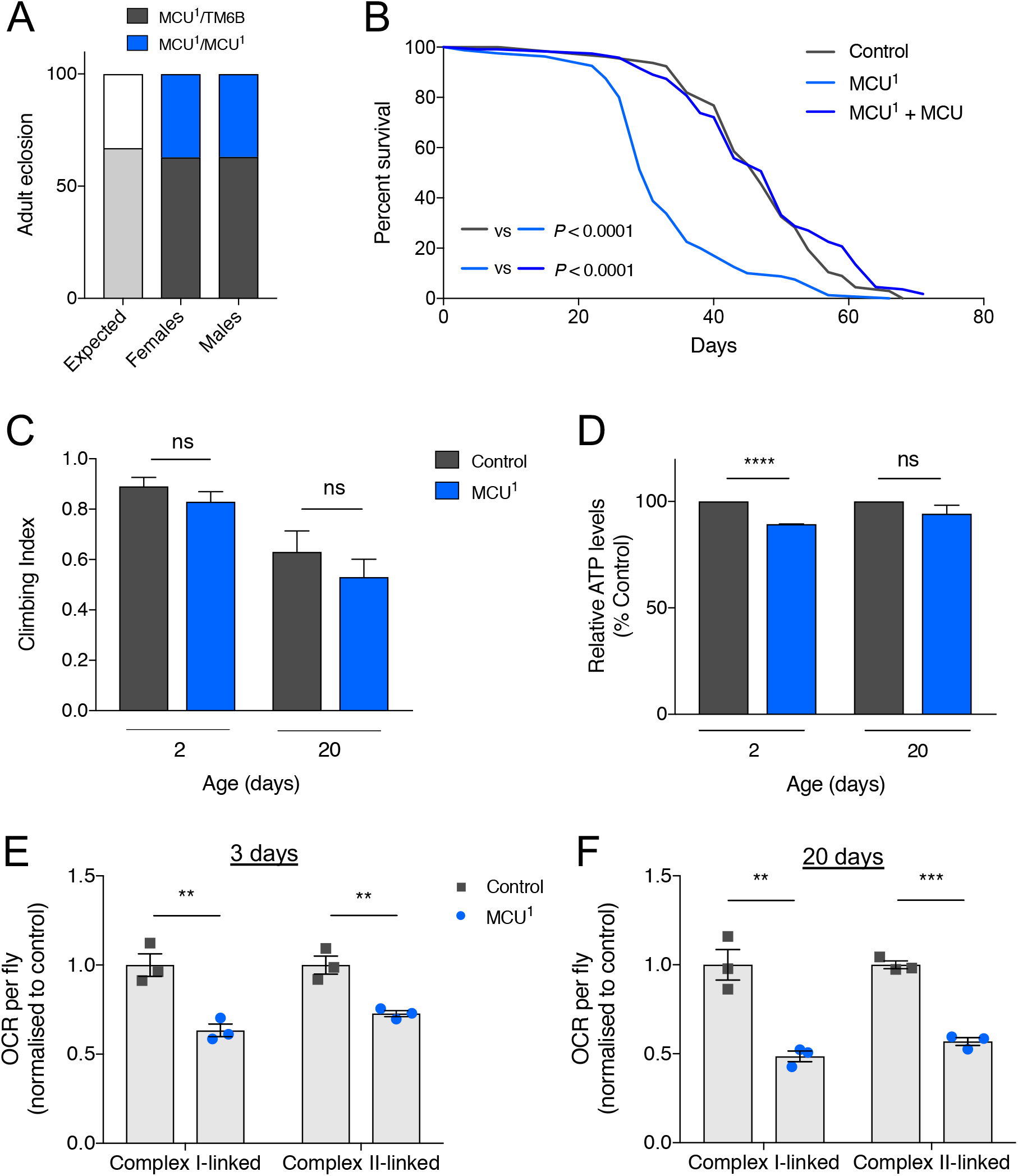
*MCU*^1^ shortens lifespan without impacting organismal phenotypes despite respiratory defects. **A.** The percentage of adult flies eclosing as homozygous *MCU*^1^ mutants versus balanced heterozygotes, together with the expected Mendelian ratio in the offspring (n > 700). **B.** Lifespan curves of *MCU*^1^ male flies compared with control (*w*^1118^) and transgenic rescue (*MCU*^1^ + *MCU*). Statistical analysis: Mantel-Cox log-rank test (n ≧ 74). **C.** Climbing assay of control (da/+) and *MCU*^1^ flies, 2 and 20 days post-eclosion. Significance measured by Kruskal-Wallis test with Dunn’s post-hoc correction for multiple comparisons (mean ± 95% confidence interval; n > 50; ns, non-significant). **D.** Relative ATP levels from control and *MCU*^1^ flies. Statistical analysis: unpaired *t*-test (mean ± SD; n 2-3; **** *p* < 0.0001, ns, non-significant). **E, F.** Oxygen consumption rate (OCR) of control and *MCU*^1^ flies at 2 (E) and 20 (F) days post-eclosion. Statistical analysis: unpaired *t*-test (mean ± SEM; n = 3; ** *P* < 0.01, *** *P* < 0.001).

We next sought to determine the effect of MCU loss on mitochondrial metabolic function. Young *MCU*^1^ mutants showed a modest but significant decrease in basal ATP levels compared to controls (Fig. 2D), which ameliorated with age. In contrast, measurement of oxygen consumption rate (OCR) revealed a marked reduction in Complex I‐ or Complex II-linked respiration in young and older flies (Fig. 2E, F). Assessing the impact on mitochondrial cell biology we found no difference in mitochondrial morphology in flight muscle (Fig. EV3A), or mitochondrial axonal transport (Fig. EV3B, C). Together these results indicate that loss of MCU affects mitochondrial respiratory capacity, but this is surprisingly well tolerated at the organismal level.

**Figure EV3.**
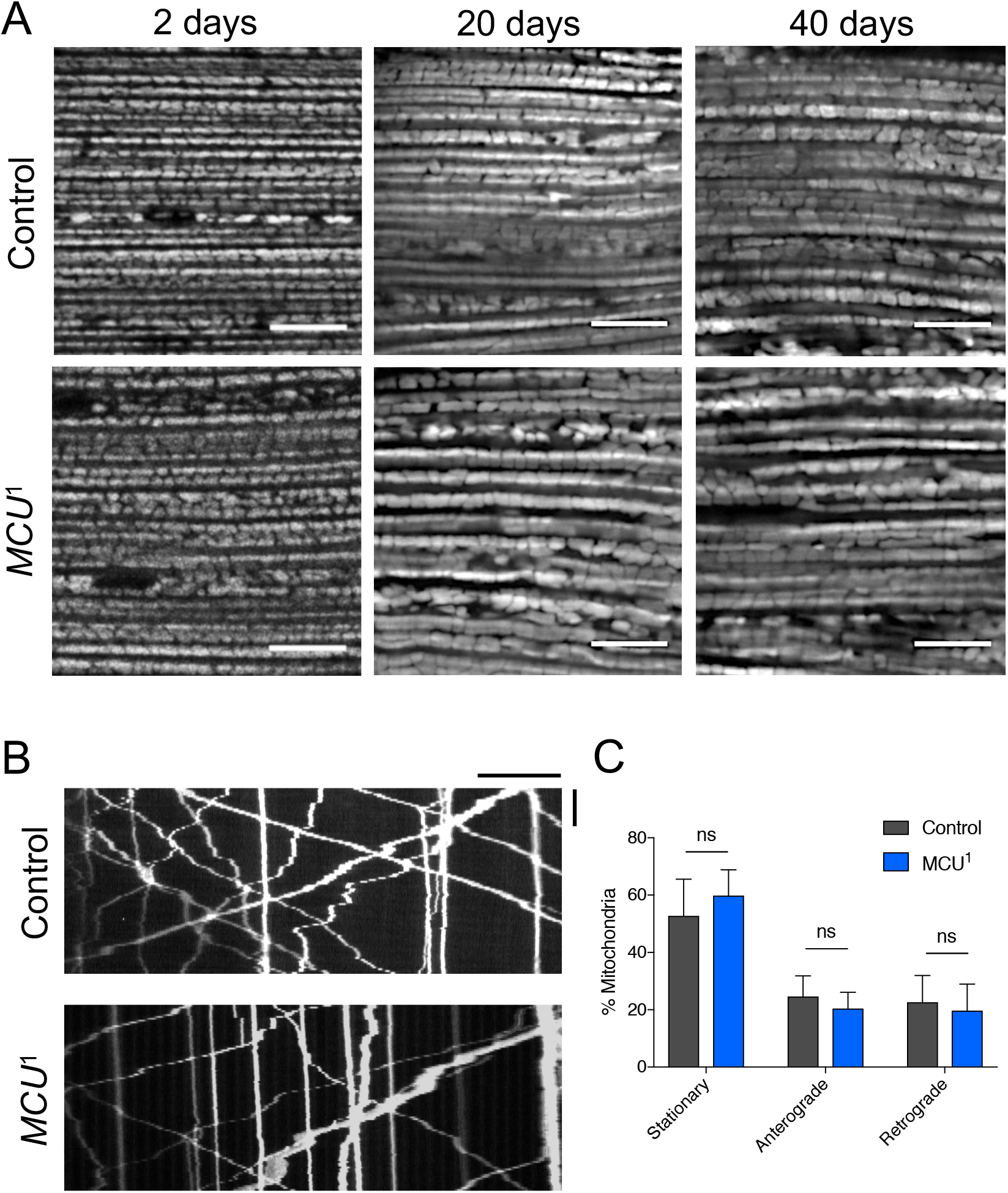
Mitochondrial morphology and dynamics are not grossly affected in *MCU*^1^ mutants. **A.** Mitochondrial morphology of control or *MCU*^1^ adult indirect flight muscle at 2, 20, or 40 days post-eclosion. Scale bar = 10 μm. Genotypes were; Control: UAS-*mito-HA-GFP*/+; *da-GAL4*/+. *MCU*^1^: UAS-*mito-HA-GFP*/+; *MCU*^1^/*MCU*^1^, *da-GAL4*. **B.** Representative kymographs of mitochondrial axonal transport in control and *MCU*^1^ larvae. Scale bars: horizontal = 20 μm, vertical = 100 s. Genotypes - Control: *CCAP-GAL4*, UAS-*mito.tdTomato*/+. *MCU*^1^: *CCAP-GAL4*, UAS-*mito.tdTomato*/+; *MCU*^1^/*MCU*^1^. **C.** Quantification of mitochondrial transport shown in B. Statistical analysis: One-way ANOVA (mean ± 95% confidence interval; n = 10; ns, non-significant).

To mutagenize *EMRE* we used a CRISPR/Cas9 based approach [20], using two simultaneously expressed transgenic guide RNAs (Fig. 3A). We isolated several indel events resulting in frameshift mutations leading to premature stop codons. Three such mutations are shown in Fig. 3B. As with *MCU*^1^, analysis of these *EMRE* mutants revealed that they all exhibit no fast mitochondrial Ca^2^+ uptake (Fig. 3C, D), in normally energised mitochondria (EV2C, D), indicating that the three mutants are functionally equivalent. We focussed on one mutant, *EMRE*^1^, which also showed a substantial reduction in the level of the mRNA transcript (Fig. 3E), for further characterisation.

**Figure 3.**
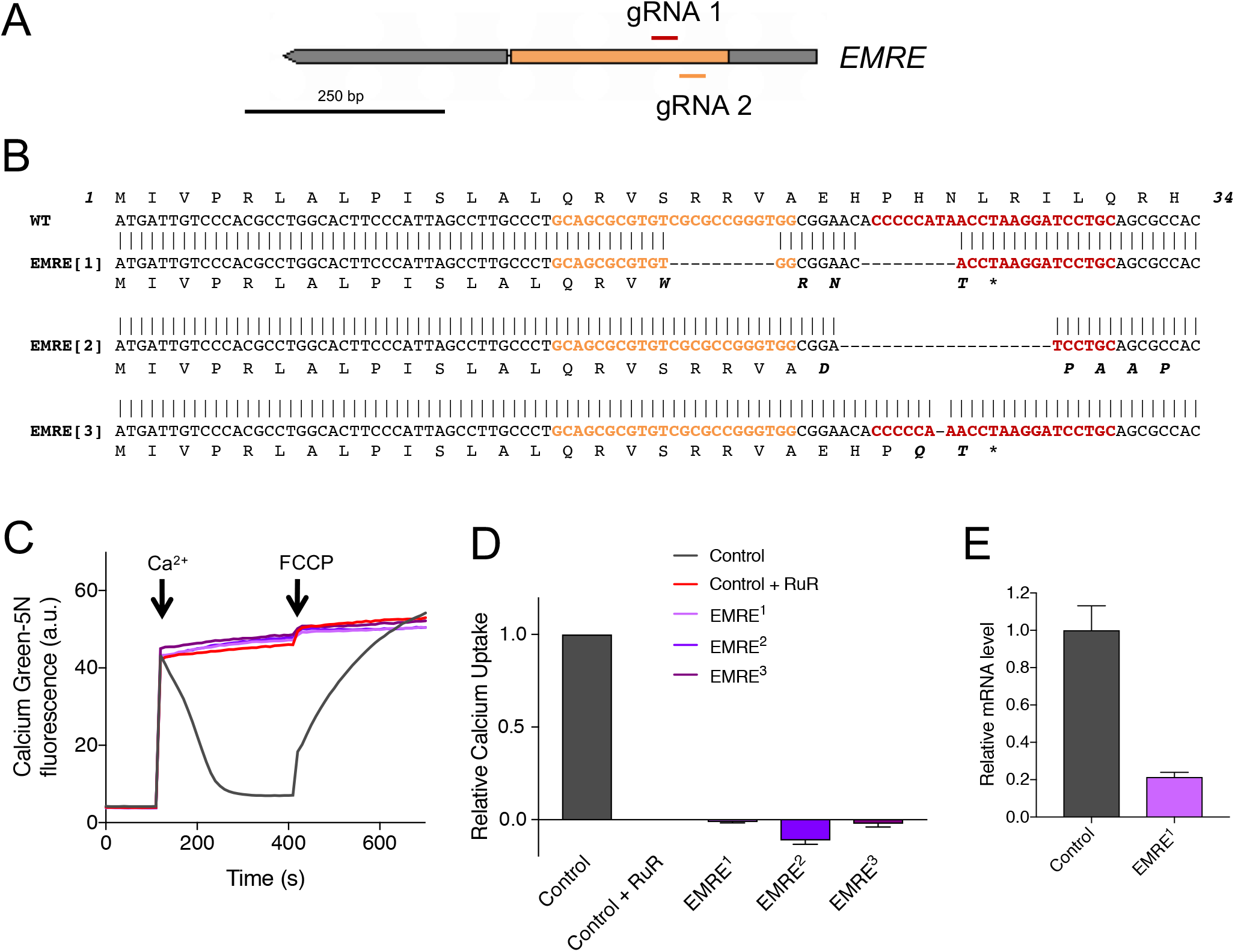
*EMRE* mutants exhibit no fast mitochondrial Ca^2+^ uptake. **A.** Overview of *EMRE* (CG17680) gene region (from FlyBase), including positions of gRNA recognition sites. **B.** Sequence alignments of wild type and *EMRE* mutants, with predicted protein sequences. **C.** Representative traces of Ca^2+^ uptake in mitochondria isolated from adult flies of the indicated genotypes after addition of 45 μM CaCl_2_. Extramitochondrial Ca^2+^ was measured by Calcium Green-5N fluorescence. Ca^2+^ was released from mitochondria by addition of 1 μM FCCP. *a.u.*: arbitrary units. The control genotype is *w*^1118^. *EMRE* mutations prevent mitochondrial Ca^2+^ uptake equivalent to the inhibitor Ruthenium Red (RuR; 2 μM). **D.** Relative uptake kinetics were determined through linear fits of Ca^2+^ uptake traces and normalized to controls (mean ± SEM; n = 3). **E.** Relative expression of *EMRE* transcript for control and *EMRE*^1^ mutants (mean ± SD; n = 3).

Similar to the *MCU*^1^ flies, *EMRE*^1^ mutants are viable, eclose at expected Mendelian ratios (Fig. 4A), and display a significantly shortened lifespan (23% reduction of median lifespan compared to control) (Fig. 4B). The climbing ability of *EMRE*^1^ mutants is slightly reduced compared to control at a young age (2 day old), but this becomes indistinguishable from control by 20 days (Fig. 4C). The basal ATP level of *EMRE*^1^ mutants is only marginally reduced at 20 days (Fig. 4D). However, in contrast to the strong reduction in respiration seen in *MCU*^1^ mutants, the Complex I‐ or Complex II-linked respiration of *EMRE*^1^ flies is either non-significant or only modestly affected compared to controls (Fig. 4E, F).

**Figure 4.**
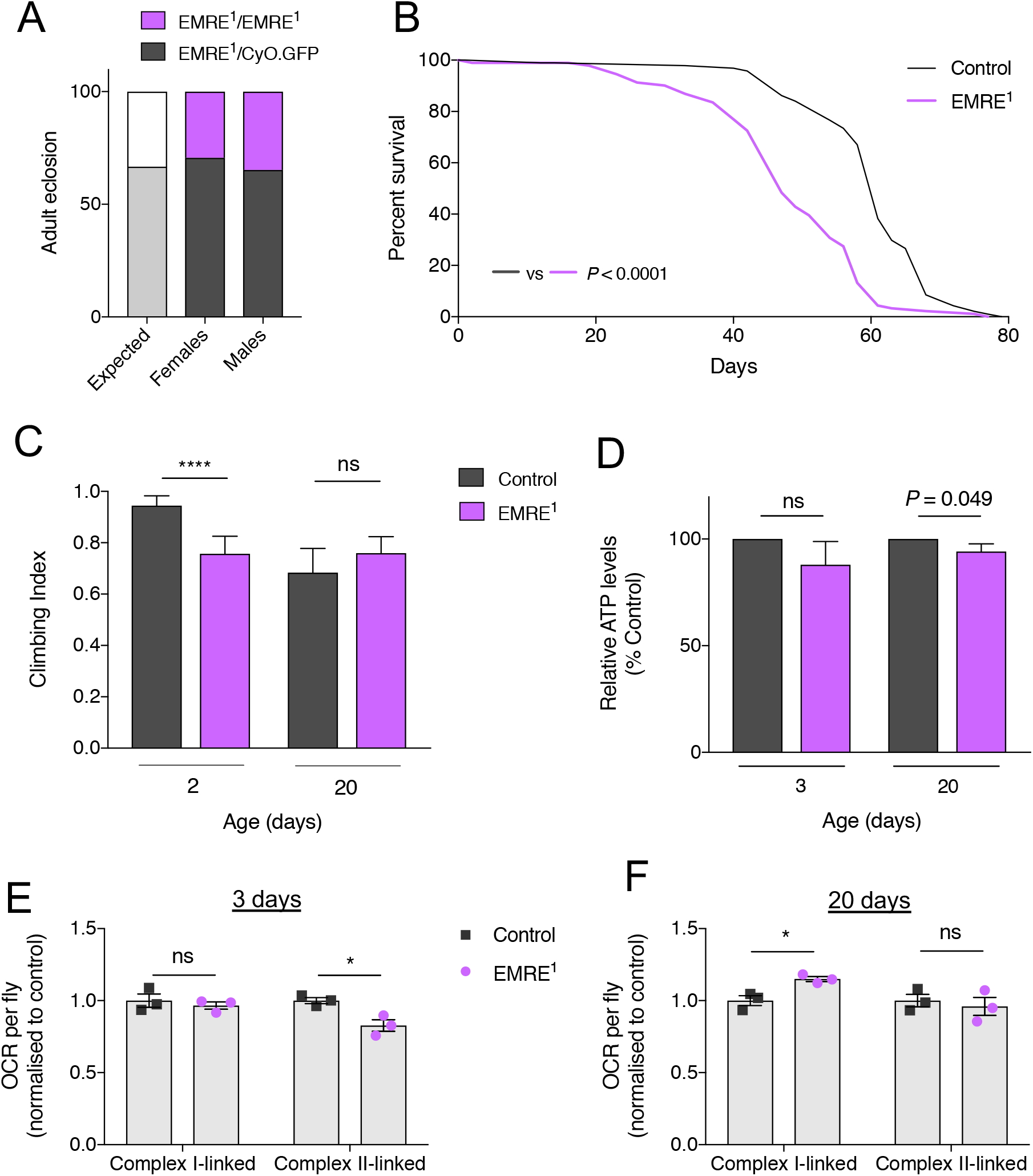
*EMRE^1^* mutants are short-lived but have very mild phenotypes. **A.** The percentage of adult flies eclosing as homozygous *EMRE*^1^ mutants versus balanced heterozygotes, together with the expected Mendelian ratio in the offspring (n > 360). **B.** Lifespan curves of *EMRE*^1^ male flies compared with control (*w*^1118^). Statistical analysis: Mantel-Coxlog-ranktest (n ≧ 91). **C.** Climbing assay of control (da/+) and *EMRE*^1^ flies, 2 and 20 days post-eclosion. Statistical analysis: Kruskal-Wallis test with Dunn’s post-hoc correction for multiple comparisons (mean ± 95% confidence interval; n > 50; **** *P* < 0.0001, ns, non-significant). **D.** Relative ATP levels from control and *EMRE^1^* flies. Statistical analysis: unpaired *t*-test (mean ± SD; n = 3; * *P* < 0.05, ns, non-significant). **E.F.** Oxygen consumption rate (OCR) of control and *EMRE*^1^ flies at 2 (panel E) and 20 (panel F) days post-eclosion. Statistical analysis: unpaired *t*-test (mean ± SEM; n = 3; * *P* < 0.05.164:390.

To target *MICU1* we again used P-element mobilisation, using MICU1^KG04119^, and isolated a large deletion spanning ~11 kb, removing half of *MICU1* and extending some 9 kb upstream of *MICU1* (Fig. 5A). This region is relatively gene-sparse and devoid of additional predicted protein coding genes. Expression analysis of homozygous *MICU1*^32^ larvae yielded no detectable transcript, establishing it as a null allele (Fig. 5B). In stark contrast to *MCU* and *EMRE* mutants, homozygous *MICU1*^32^ mutants are larval lethal with a few animals reaching the third instar stage (Fig. 5C). Supporting this, ubiquitous expression of two independent RNAi transgenes also caused developmental lethality (data not shown). Importantly, full viability to adult stage was restored upon ubiquitous expression of HA-tagged MICU1, by either the A or B isoforms (Fig. 5D). Moreover, organismal vitality, as measured by climbing ability, was also completely restored (Fig. 5E).

**Figure 5.**
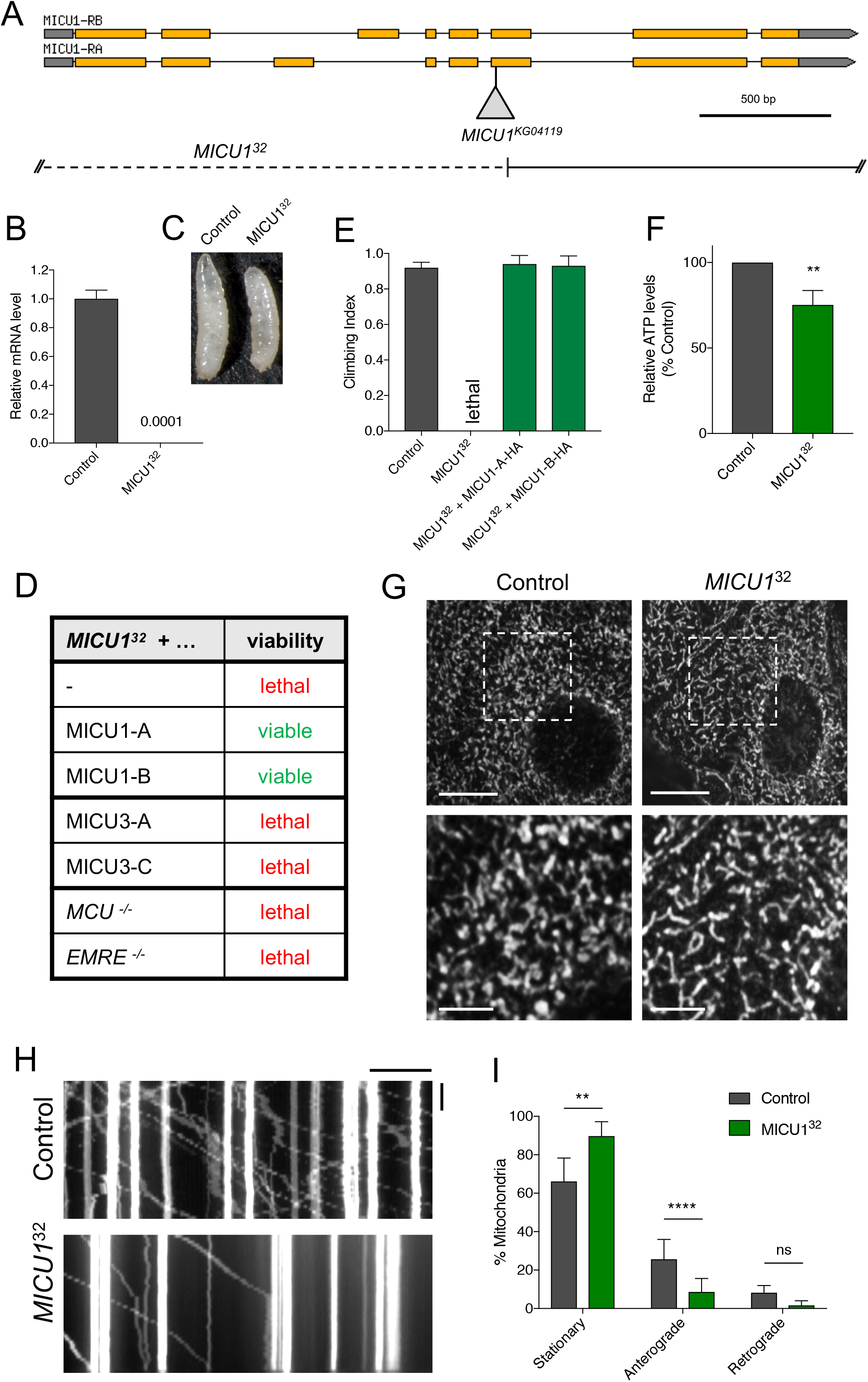
*MICU1*^32^ mutants are lethal, have reduced ATP and mitochondrial transport, and are not rescued by *MCU*^1^ or *EMRE*^1^. **A.** Overview of *MICU1* (CG4495) gene region (from FlyBase). The P{SUPor-P}MICU1^KG04119^ transposable element used to generate *MICU1*^32^ is displayed. **B.** Relative expression of *MICU1* transcript for control and *MICU1*^32^ larvae (mean ± SD; n = 3). **C.** Control and *MICU1*^32^ larvae at lethal phase. **D.** Table of viability of *MICU1*^32^ rescue by transgenic expression of *MICU1* or *MICU3* isoforms, or loss of *MCU* or *EMRE*. Transgenic expression was induced using ubiquitous drivers; *arm-GAL4* for *MICU1* and *da-GAL4* for *MICU3*. **E.** Climbing assay of control flies and *MICU1*^32^ mutants with ubiquitous (*arm-GAL4*) driven transgenic re-expression of HA-tagged MICU1-A and တB isoforms. Statistical analysis: Kruskal-Wallis test with Dunn’s post-hoc correction for multiple comparisons (mean ± 95% confidence interval; n ≧ 40; **** *P* < 0.0001). **F.** Relative ATP levels from control and *MICU1*^32^ larvae. Statistical analysis: unpaired *t*-test (mean ± SD; n = 4; ** *P=* < 0.0011). **G.** Confocal microscopy analysis of epidermal cells in control and *MICU1*^32^ larvae immunostained with the mitochondrial marker anti-ATP5A. Boxed areas are enlarged below. Scale bars are 10 μm (top) and 4 μm (bottom). **H.** Representative kymographs of mitochondrial axonal transport in control and *MICU1*^32^ larvae. Scale bars: horizontal = 20 μm, vertical = 50 s. Genotypes - Control: *M12-GAL4*, UAS-*mito-HA-GFP/+. MICU1^32^: MICU1^32^/MICU1^32^; M12-GAL4*, UAS-*mito-HA-GFP*/+. **I.** Quantification of mitochondrial transport shown in E. Statistical analysis: One-way ANOVA (mean ± 95% confidence interval; n = 6 (control) and 11 (mutant); ** *P* < 0.01, **** *P* < 0.0001, ns, non-significant).

The severely reduced viability of *MICU1*^32^ mutants precluded the possibility of performing respirometry, but analysing ATP levels we found a significant reduction (Fig. 5F), indicative of a substantial mitochondrial impairment. This prompted us to investigate other aspects of mitochondrial homeostasis. Assessing mitochondrial morphology in larval epidermal cells we found that the distribution of mitochondria appeared altered, less inter-connected and more diffuse (Fig. 5G). We also found that axonal transport of mitochondria is impaired in *MICU1*^32^ larvae (Fig. 5H, I). Collectively, these data show the presence of multiple mitochondrial defects that together might be responsible for lethality of the *MICU1*^32^ mutants.

*Drosophila* do not have an orthologue of the *MICU1* paralogue, *MICU2*, but *MICU3* is conserved, encoded by *CG4662* (Fig. 6A). Interestingly, *Drosophila MICU3* also appears to be neuronally restricted (see FlyBase and [21]), as in mammals [9]. Currently, very little is known about the function of MICU3, and no *in vivo* studies have been reported. To assess its role *in vivo* we used CRISPR/Cas9 to induce indel mutations. One of these mutations, *MICU3*^27^, a single base deletion (Fig. 6B, C), which abolishes a *MboII* restriction site (Fig. 6D), leads to a frameshift and early truncation. This mutation also substantially de-stabilises the *MICU3* transcript (Fig. 6E). In striking contrast to loss of *MICU1*, homozygous *MICU3*^27^ mutants are fully viable (Fig. 6F), though lifespan was modestly (7% reduction in median lifespan) but significantly reduced (Fig. 6G). In addition, these mutants exhibited a significant climbing defect in young and older flies (Fig. 6H). These results indicate a function for MICU3 in proper maintenance of neuronal function, consistent with its neuronally-restricted expression. However, analysing mitochondrial respiration from heads of *MICU3*^27^ mutants, we observed no significant differences compared to control (Fig. 6I).

**Figure 6.**
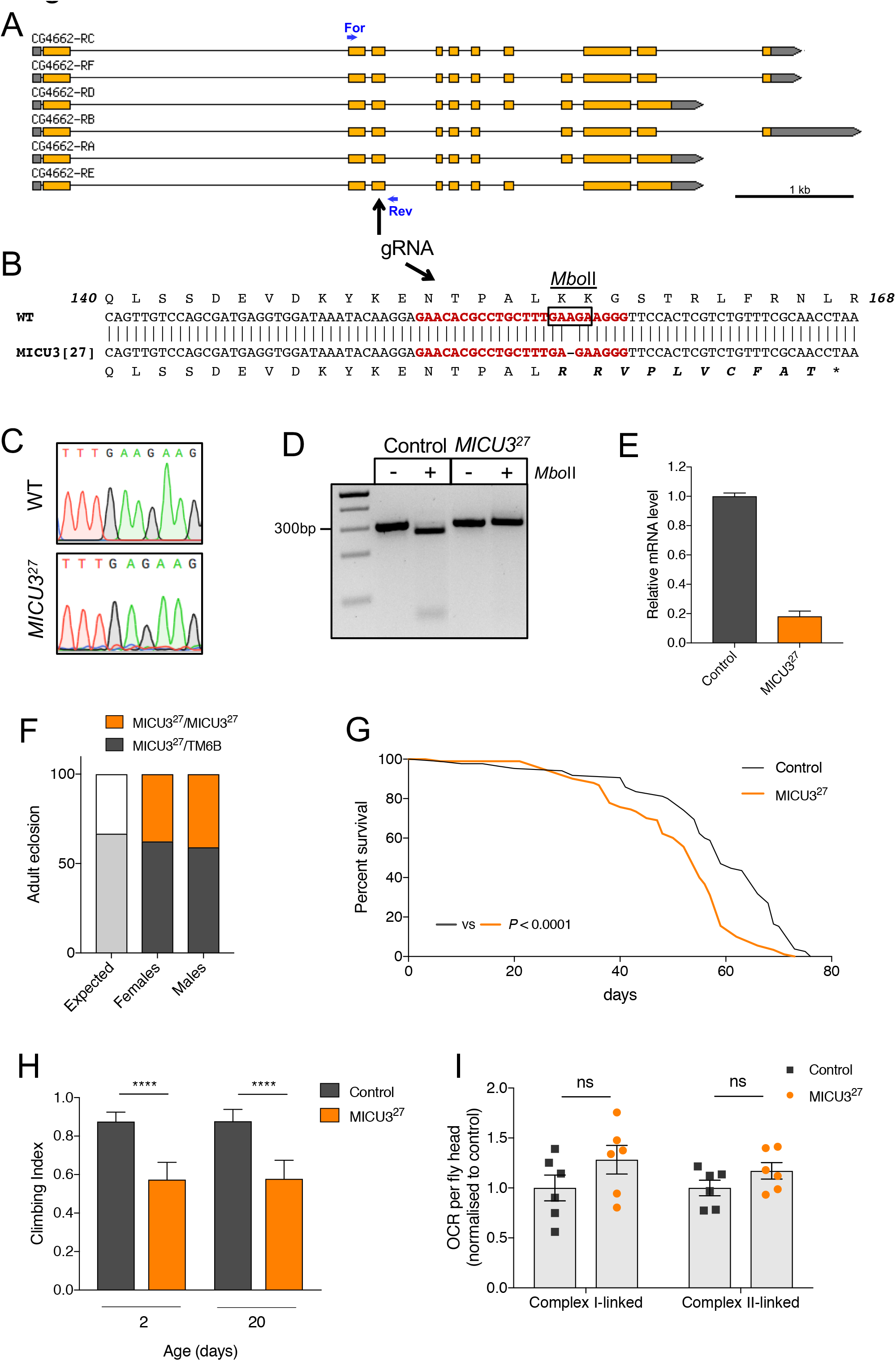
*MICU3* loss-of-function is largely benign. **A.** Overview of *MICU3* (CG4662) generegion (from FlyBase), including positions of gRNA recognition site and for ward/reverse primers used for Restriction Fragment Length Polymorphism (RFLP) analysis used in D. **B.** Sequence alignment of wild type and *MICU3*^27^, with relative predicted protein sequences. The gRNA recognition site is highlighted in red. Box denotes *Mbo* II cleavage site. **C.** Representative sequencing chromatograms for wild-type and *MICU3*^27^ showing the single base deletion of *MICU3*^27^. **D.** RFLP analysis of control and *MICU3*^27^ PCR products with *Mbo* II. **E.** Relative expression of *MICU3* transcript for control and *MICU3*^27^ flies (mean ± SD; n = 3). **F.** The percentage of adult flies eclosing as homozygous *MICU3*^27^ mutants versus balanced heterozygotes, together with the expected Mendelian ratio in the offspring (n > 225). **G.** Lifespan curves of *MICU3*^27^ male flies compared with control (*w*^1118^). Statistical analysis: Mantel-Cox log-rank test (n ≧ 84). **H.** Climbing assay of control (da/+) and *MICU3*^27^ flies, 2 and 20 days post-eclosion. Statistical analysis: Kruskal-Wallis test with Dunn’s post-hoc correction for multiple comparisons (mean ± 95% confidence interval; n > 50; **** *P* < 0.0001). **I.** Oxygen consumption rate (OCR) of control and *MICU3*^27^ flies at 2 days post-eclosion. Statistical analysis: unpaired *t*-test (mean ± SEM; n = 3).

In order to investigate the functional relationship between the various uniporter components we undertook a number of genetic interaction studies. First, since *Drosophila* MICU1 and MICU3 share a fair degree of homology (~49% similarity, ~31% identity between MICU1-B and MICU3-C), we reasoned that they may share some functional overlap. To address this, we asked whether normally neuronally-restricted MICU3 could functionally substitute for MICU1. Thus, we ectopically expressed *MICU3* ubiquitously in *MICU1*^32^ mutants. Here we chose to express isoforms A and C, as these cover all of the predicted coding regions (Fig. 6A). However, neither MICU3 isoform was able to restore viability of *MICU1*^32^ mutants or shift the lethal phase (Fig. 5H), indicating that MICU3 is not functionally equivalent to MICU1 *in vivo*.

MICU1 has been shown to provide gatekeeper function for the uniporter channel with loss of MICU1 causing unregulated mitochondrial Ca^2+^ uptake. It has also been shown that in mice lacking *MICU1*, genetic reduction of *EMRE* substantially ameliorates the *MICU1* phenotypes [19]. Thus, we reasoned that the lethality of the *MICU1*^32^ mutants is caused by unregulated Ca^2+^ entry, which should be prevented by loss of the MCU channel. To test this, we combined homozygous *MICU1*^32^ and *MCU*^1^ mutants and, to our surprise, found that this did not suppress the lethality or noticeably shift the lethal phase (Fig. 5H). We corroborated this finding by combining *MICU1*^32^ mutants with *EMRE*^1^ mutants, with the same result (Fig. 5H).

Overexpression paradigms disrupting uniporter stoichiometry have previously been used to interrogate the functional relationships of uniporter components [22]. Using a classic eye morphology assay as a read-out of the impact of genetic interactions on cell and tissue viability, we first found that overexpression of any of the uniporter components alone in the eye, using a *GMR-GAL4* driver, had no effect on eye or ommatidial morphology (Fig. EV4). This indicates that overabundance of any uniporter component, including MCU itself, is insufficient to grossly disrupt mitochondrial Ca^2+^ homeostasis as expected. However, the co-expression of *MCU* and *EMRE* caused a dramatic disruption of eye morphology with a general loss of retinal pigment and ommatidia resulting in a glazed appearance with occasional black, necrotic patches (Fig. 7A). This effect is consistent with the cooperative actions of MCU and EMRE to create the channel, and in line with reconstitution experiments in yeast showing that expression of mammalian *MCU* and *EMRE* are necessary and sufficient to elicit Ca^2+^ uniporter activity. The gross disruption of eye integrity also demonstrates the catastrophic effects of unregulated mitochondrial Ca^2+^ entry.

**Figure EV4.**
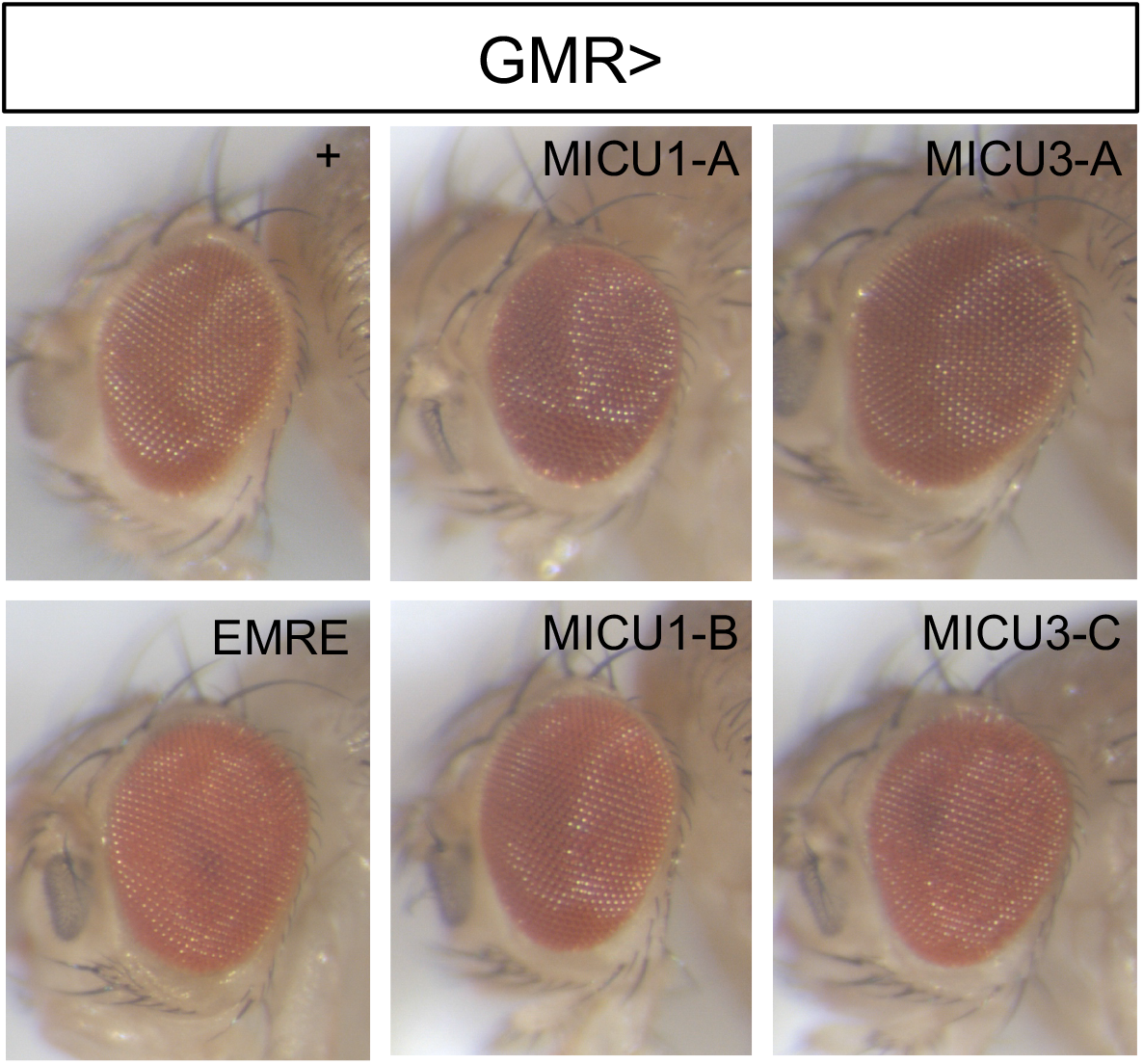
Eye-driven overexpression of individual uniporter components is benign. Representative images of eye morphology for GMR-driven expression of individual uniporter transgenes, or *GMR-GAL4*/+ control.

**Figure 7.**
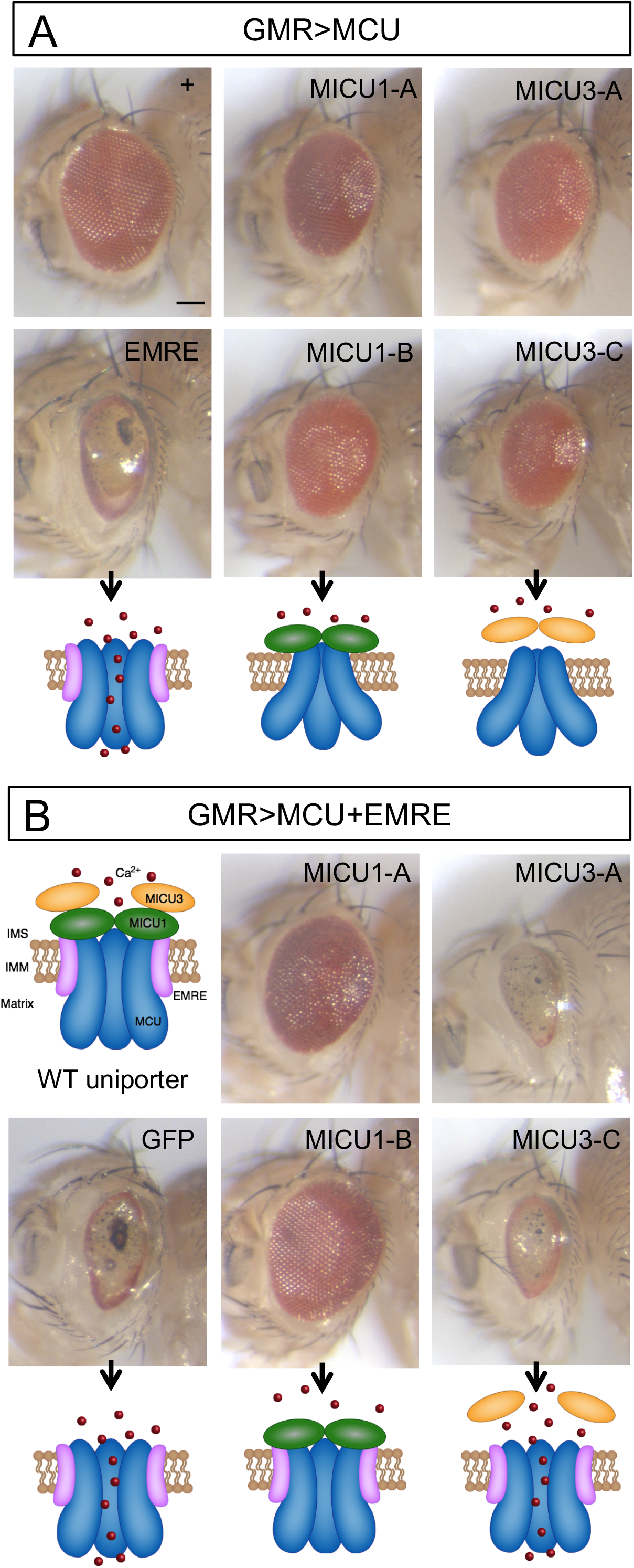
Genetic interactions of overexpression uniporter components. **A.** A line bearing the eye-specific *GMR-GAL4* driver and *UAS-MCU* transgene was crossed to transgenes of the indicated uniporter components, or to a control (+) *w*^1118^ line. **B.** A line bearing a combination of the eye-specific *GMR-GAL4* driver and both a *UAS-MCU* and *UAS-EMRE* transgenes was crossed to transgenes of the indicated components, or to a control (GFP) transgene. Below the micrograph images are schematic cartoons of the proposed status of the uniporter based on the composition of the overexpressed components. Scale bar = 100 μm.

In contrast, co-expression of *MCU* with either *MICU1* or *MICU3* was not so detrimental, although in all cases it caused a very mild disruption of the ommatidial arrangement resulting in a mild ‘roughened’ appearance (Fig. 7A). This system allowed us to test possible functional differences between different isoforms of *MICU1* and *MICU3*. In general, the co-expression of all isoforms with *MCU* caused similar effects, though it is notable that the expression of *MICU3-C* gave a slightly stronger phenotype, that may reflect its greater level of expression compared to *MICU3-A* (Fig. EV1F). The enhanced phenotype of *MCU:MICU3* co-expression is consistent with a recent report that MICU3 enhances MCU-mediated Ca^2+^ uptake [9].

We next reasoned that if the dramatic eye phenotype caused by *MCU* and *EMRE* co-expression was due to extra unregulated channels and excessive Ca^2+^ uptake, this may be ameliorated by co-expression of the *MICU1* gatekeeper. Remarkably, co-expression of either *MICU1-A* or *-B* with *MCU* and *EMRE* prevented the *MCU:EMRE* phenotype (Fig. 7B). Co-expression of a *GFP* control confirmed that this was not due to titration of multiple UAS lines. Notably, while both isoforms provided excellent rescue, isoform B appeared slightly more effective than isoform A. These observations are entirely consistent with MICU1 providing the main gatekeeper function for the uniporter.

We also used this assay to test the functionality of MICU3 in this context. In striking contrast to *MICU1* expression, co-expression of either *MICU3-A* or *-C* with *MCU* and *EMRE* provided no suppressing effect for the *MCU:EMRE* phenotype (Fig. 7B). These results are consistent with MICU3 not being a major uniporter gatekeeper *in vivo*.

## Discussion

The capacity of mitochondria to take up Ca^2+^ has important implications for cellular homeostasis as it regulates fundamental processes from metabolism to cell death. In this context, the mitochondrial Ca^2+^ uniporter plays a crucial function, driving the rapid entry of Ca^2+^ into mitochondria. In order to better understand the physiological role of mitochondrial Ca^2+^ uptake, we have used the genetically powerful model *Drosophila* to manipulate the conserved set of dedicated channel and regulatory proteins that form the complex.

The fact that *MCU* mutants are viable and fertile with no gross morphological or behavioural defects was initially surprising given the historical importance of mitochondrial Ca^2+^, but corroborates another recent report of fly *MCU* mutants [22], and is consistent with studies in mice and worms where deletion of the *MCU* orthologues are essentially benign at the organismal level under basal conditions [15,23]. Importantly, however, fly *MCU* mutants are significantly shorter-lived than controls. This situation is mirrored by *EMRE* mutants, albeit with a smaller impact on lifespan. The reason for the shortened lifespans is currently unknown but may reflect the effects of a chronic bioenergetic deficit evident from the OCR measurements. Accordingly, *MCU* mutants show a greater respiration defect compared to *EMRE* mutants, consistent with their respective impacts on lifespan. The respiratory impairment could be due to the previously reported increase in oxidative stress that occurs in the *MCU* mutants [22], which has yet to be assessed in *EMRE* mutants. Alternatively, the short lifespan may be due to a myriad of potential metabolic imbalances, such as disruption of NADH/NAD+ levels. Chronic adaptations may also occur through transcriptional responses. Further studies analysing the metabolic and transcriptional changes occurring in these flies will shed light on this fundamental question.

Nevertheless, the fact that the *EMRE* mutants are also relatively benign at the organismal level corroborates the surprising viability of *MCU* mutants. Considering this, it is striking that flies, as in mice and worms, consistently show an ability to compensate for the lack of fast mitochondrial Ca^2^+ uptake, suggesting the induction of some adaptive mechanism as discussed by others [16,24]. While alternative routes of mitochondrial Ca^2+^ entry must exist, since matrix Ca^2+^ is not abolished in *MCU* KO mice [15], proposed mechanisms are speculative, and it is unclear whether they constitute a compensatory adaptation for fast Ca^2^+ uptake or simply allow gradual, slow accumulation [24]. On the other hand, rapid mitochondrial Ca^2+^ uptake mediated by MCU is thought to constitute a specific metabolic regulatory mechanism, e.g. to increase ATP production, under certain conditions such as strenuous exercise or pathological conditions [25], which is partly evident in the *MCU* KO mice [15] or heart-specific conditional KO [17]. Such important physiological roles would not necessarily be apparent under basal conditions in flies. Indeed, MCU has also been proposed to promote wound healing [23], however, our preliminary studies did not find evidence supporting this. The current study presents a summary of the requirements of uniporter components under basal conditions, and further work will be needed to evaluate the role of the uniporter in the full range of physiological conditions.

In seeking to understand the importance of the regulatory components of the uniporter we also developed loss-of-function models for *MICU1* and *MICU3*. In striking contrast to *MCU* and *EMRE* mutants, loss of *MICU1* results in larval lethality which is associated with alterations in mitochondrial morphology and motility, and a reduced level of total ATP. In line with its role as the principle gatekeeper of the uniporter and the knowledge that excess mitochondrial Ca^2+^ triggers cell death, we reasoned that the lethality was due to Ca^2+^ accumulation in the mitochondrial matrix through unregulated MCU-EMRE channels. Supporting this, we observed that dual overexpression of *MCU* and *EMRE* in the eye leads to substantial loss of retinal tissue; concomitant overexpression of *MICU1* is sufficient to prevent this phenotype, consistent with MICU1 re-establishing appropriately regulated uniporter channels.

However, one striking observation that was most surprising to us was the inability of *MCU* or *EMRE* mutants to rescue the *MICU1* mutant lethality. This result is particularly puzzling since it was recently shown that mice lacking *MICU1*, which present multiple pathogenic phenotypes, are substantially rescued by genetic reduction of *EMRE* levels [19]. While the reason for the lack of rescue in flies is unclear, we postulate that this suggests the function of MICU1 is not limited to uniporter-dependent Ca^2+^ uptake. We do not currently know whether the lethality of *MICU1* mutants is specifically due to excessive mitochondrial Ca^2+^ levels, however, it appears to be independent of fast mitochondrial Ca^2+^ uptake as this is eliminated in *MCU* and *EMRE* mutants. As noted above, other routes of Ca^2+^ uptake into mitochondrial clearly exist but the mechanisms that regulate them are uncertain. These *Drosophila* models are ideally suited for unbiased genetic screening to uncover such fundamental regulatory mechanisms.

In contrast to *MICU1*, loss of *MICU3* was overall well tolerated at the organismal level. Functional analysis of MICU3 is to date extremely limited, but the neuronally-restricted expression led us to anticipate that these mutants might have more neurological-specific phenotypes, which was at least partly borne out. Whereas longevity of these mutants was only minimally affected, they exhibited a notable locomotor deficit even in young flies. We initially hypothesized that MICU3 may be able to act redundantly with MICU1 but attempts to transgenically rescue *MICU1* mutants by ectopic *MICU3* expression were unsuccessful. This result is consistent with a recent report showing that MICU3 binds to MICU1 but apparently enhances mitochondrial Ca^2+^ uptake [9].

In summary, we present a comprehensive analysis of the conserved components of the mitochondrial Ca^2+^ importer and its regulators. While loss of the various components results in dramatically different organismal phenotypes, ranging from the most severe deficit exemplified by the *MICU1* mutants to the very mild consequences of mutating *MICU3*, such diverse phenotypes mirror the situation reported in humans so far. The first described patients with *MICU1* mutations exhibit a severe, complex neurological condition accompanied by muscular dystrophy and congenital myopathy, clearly associated with mitochondrial dysfunction [26], whereas a later study reported *MICU1* patients with a relatively mild fatigue syndrome [27]. One explanation for the reported phenotypic variability is that the consequence of perturbing mitochondrial Ca^2+^ uptake can be influenced by additional factors, the most obvious being genetic background. The genetic tools described here open up the possibility for a thorough analysis of the uniporter function in a powerful genetic model organism which will advance our understanding of the role of mitochondrial Ca^2+^ in health and disease.

## Materials and Methods

### *Drosophila* husbandry, stocks and mutagenesis

Flies were raised under standard conditions at 25 °C on food consisting of agar, cornmeal, molasses, propionic acid and yeast in a 12h:12h light:dark cycle. The following strains were obtained from the Bloomington *Drosophila* Stock Center (RRID:SCR_006457): *w*^1118^ (RRID:BDSC_6326), MCU^EY01803^ (RRID:BDSC_16357), MICU1^KG04119^ (RRID:BDSC_13588), *da-GAL4* (RRID:BDSC_55850), *arm-GAL4* (RRID:BDSC_1560), *GMR-GAL4* (RRID:BDSC_1104), *Mef2-GAL4* (RRID:BDSC_27390), *CCAP-GAL4* (RRID:BDSC_25685), *M12-GAL4* (RRID:BDSC_2702), *UAS-mito-HA-GFP* (RRID:BDSC_8442, RRID:BDSC_8443), *UAS-mito.tdTomato* (RRID:DGGR_117015). UAS-*MICU1-A-HA* (F000962) was obtained from the FlyORF collection [28].

Mobilisation of the *MCU*^EY01803^ and *MICU1*^KG04119^ transposable elements was used to generate *MCU* (*CG18769*) and *MICU1* (*CG4495*) mutants, termed *MCU*^1^ and *MICU1*^32^. For *EMRE* (*CG17680*) and *MICU3* (*CG4662*), a CRISPR-based strategy was employed. Here, gRNAs were evaluated using an online tool (http://tools.flycrispr.molbio.wisc.edu/targetFinder/) [29]. Where possible, gRNAs towards the 5’ end of the protein coding sequence without predicted off-targets were selected. gRNAs were cloned into the pCFD4 vector (Addgene 49411), and the resulting constructs were verified by sequencing before being sent for transgenesis by phiC31 site-directed integration into attP40 and attP2 sites (BestGene Inc., or Dept. of Genetics, University of Cambridge). These gRNA-expressing flies were crossed to *y*^1^ M{nos-cas9, w^+^}ZH-2A w* (RRID: BDSC_54591) for mutagenesis. Mutant lines were screened via sequencing of PCR products spanning the targeted gene region. Forward (F) and reverse (R) primer sequences used for DNA analysis of *MCU*^1^ shown in Figure 1B were: F (5’-3’), GCAACTTCAGCATATGACC and R (5’-3’), GGAATTGGGATGCCATAGC. The mapped breakpoints for each mutant are as follows: *MCU*^1^, 3L:6550718-6552274; *EMRE*^1^, 2R:18147830-18147821, 2R:18147812-18147804; *EMRE*^2^, 2R: 18147814-18147795; *EMRE*^3^, 2R:18147805-18147805; MICU1^32^, 2L:7173086-7184065; *MICU3*^27^, 3R: 19850278-19850278. All genomic coordinates are according to FlyBase/BDGP Release 6 [30]. All the mutants lines used in this study were back-crossed to an isogenic *w*^1118^ strain (RRID:BDSC_6326), for 4-6 generations before use.

### Generation of transgenic lines

The following transgenes were generated by cloning into the pUAST-attB vector (BestGene Inc.). *UAS-MCU: MCU* was amplified from cDNA clone LD26402 (equivalent to isoforms A, B, C and D) and inserted between the *Not* I and *Xho* I sites. *UAS-MICU1-B-HA*: *MICU1-B* was amplified from cDNA IP17639 to include a single 3’ HA tag, and was inserted between the *Not* I and *Xba* I sites. *UAS-EMRE-myc: EMRE* was amplified from genomic DNA with primers encoding a single 3’ Myc tag, and was inserted between the *EcoR* I and *Xho* I sites. *UAS-MICU3-A-V5, UAS-MICU3-C-V5: MICU3-A* and *MICU3-C* were amplified from cDNA clones ID23951 and RH09265 respectively, including a single 3’ V5 tag, and were inserted between the *Not* I and *Xba* I sites.

All cDNA clones were obtained from *Drosophila* Genomics Resource Center (Bloomington, Indiana). Constructs were verified by sequencing before being sent for transgenesis by phiC31 site-directed integration into attP40 and attP2 sites (BestGene Inc., or Dept. of Genetics, University of Cambridge). For all integration events, multiple independent lines were initially isolated, verified by PCR and assessed for consistent effects before selecting a single line of each integration site for further study.

### Mitochondrial membrane potential and calcium flux assays

Mitochondria were prepared from ~50 whole adult flies by differential centrifugation. Samples were homogenized with a Dounce glass potter and a loose-fitting glass pestle in a mannitol-sucrose buffer (225 mM mannitol, 75 mM sucrose, 5 mM HEPES, 0.1 mM EGTA, pH 7.4) supplemented with 2% BSA. Samples were centrifuged at 1,500 × *g* at 4 °C for 6 min. The supernatant was filtered through a fine mesh, and centrifuged at 7,000 × *g* at 4 °C for 6 min. The resulting pellet was resuspended in mannitol-sucrose buffer without BSA before being centrifuged at 7,000 x *g* under the same conditions as above and resuspended in a small volume (~50 μL) of mannitol-sucrose buffer. Protein concentration was measured using the Biuret test.

Mitochondrial membrane potential of isolated mitochondria was measured based on the fluorescence quenching of Rhodamine123 (Rh123; Molecular Probes) and mitochondrial Ca^2^+ fluxes were measured by Calcium Green 5N (Molecular Probes) fluorescence at 25 °C [31] using a Fluoroskan Ascent FL(Thermo Electron) plate reader (excitation and emission wavelengths of 485 and 538 nm, respectively with a 10 nm bandpass filter) at a mitochondrial concentration of 1 mg/mL. The incubation medium contained 250 mM sucrose, 10 mM MOPS-Tris, 5 mM/2.5 mM glutamate/malate-Tris, 5 mM Pi-Tris, 10 μM EGTA, and 0.4 μM Rhodamine123, or 0.5 μM Calcium Green 5N, pH 7.4. Addition of Ruthenium Red (RuR, 2 μM) was made directly into the well containing the assay medium before mitochondria were added. Further additions were made as indicated in the figure legends.

### Locomotor and lifespan assays

Climbing (negative geotaxis assay) was assessed as previously described with minor modifications [32]. Briefly, for climbing, 20-25 males were placed into the first chamber of a ‘Benzer’ counter-current apparatus, tapped to the bottom, and given 10 s to climb a 10 cm distance. This procedure was repeated five times, and the number of flies remaining in each chamber was counted. The weighted performance of several group of flies for each genotype was normalized to the maximum possible score and expressed as Climbing Index.

For lifespan experiments, flies were grown under identical conditions at low-density. Groups of approximately 20-25 males of each genotype were collected under very light anaesthesia, placed into separate vials with food and maintained at 25 °C. Flies were transferred into vials containing fresh food every 2-3 days, and the number of dead flies was recorded. Percent survival was calculated at the end of the experiment after correcting for any loss resulting from handling.

### Mitochondrial protein enrichment

Crude mitochondrial extracts were obtained from approximately 100 flies per sample. After 5 min on ice, flies were homogenized in a glass tissue grinder containing 2 mL of cold mitochondrial isotonic buffer (225 mM mannitol, 75 mM sucrose, 5 mM HEPES, 0.5 mM EGTA, 2 mM taurine, pH 7.25) for 30-60 seconds until uniform. Subsequently, the homogenates were centrifuged at 500 × *g* at 4 °C for 5 min. The resulting supernatants were passed through a 100 μm nylon sieve (Cell Strainer REF 352360, BD Falcon, USA), centrifuged at 11,000 × *g* for 10 min at 4 °C, and pellets were stored at - 80 °C until use. The final mitochondrial pellets were subsequently used for lysis in 50 μL of RIPA buffer (50 mM Tris-HCl, pH 8.0; 150 mM NaCl; 1 mM EDTA, 0.5% SDS, 1% (vol/vol) Triton X-100) with cOmplete mini EDTA-free protease inhibitors (Roche) for 10 min on ice. After carrying out three freeze-thaw cycles with dry ice and a 37 °C water bath, the lysates were centrifuged at 20,000 × *g* for 5 min and the supernatants taken for SDS page.

### Immunoblotting

For MCU and EMRE expression analysis, mitochondrial proteins were isolated from whole adult flies according to the method described in the previous section. For MICU1 and MICU3 overexpression analysis flies were homogenized in a PBS-based lysis buffer with lithium dodecyl sulfate containing β-Mercaptoethanol and supplemented with cOmplete mini EDTA-free protease inhibitors (Roche). Equivalent amounts of proteins were resolved by SDS-PAGE and transferred onto nitrocellulose membrane using a semi-dry Transblot apparatus (BioRad) according to the manufacturer’s instructions. The membranes were blocked in TBST (0.15 M NaCl and 10 mM Tris-HCl; pH 7.5, 0.1% Tween 20) containing 5% (w/v) dried non-fat milk (blocking solution) for 1 h at room temperature and probed with the indicated primary antibody before being incubated with the appropriate HRP-conjugated secondary antibody. Antibody complexes were visualized by an ECL-Plus enhanced chemiluminescence detection kit (Amersham) using a ChemiDoc XRS+ molecular imager (BioRad).

### Antibodies

For immunoblot experiments, the following antibodies were used: mouse anti-ATP5A (Abcam ab14748; RRID:AB_301447; 1:20000), rabbit anti-HA (Abcam ab9110; RRID:AB_307019; 1:1000), mouse anti-α-Tubulin (Sigma T9026, clone DM1A; RRID:AB_477593; 1:1500), rabbit anti-Porin (Millipore PC548; RRID:AB_2257155; 1:5000), anti-V5 (Thermo Fisher Scientific R960-25; RRID:AB_2556564; 1:2000), mouse anti-Myc tag (Cell Signaling, clone 9B11; RRID:AB_331783; 1:800). Horseradish peroxidase-conjugated secondary antibodies: anti-mouse (Abcam ab6789-1; RRID:AB_955439; 1:5000-1:40000), anti-rabbit (Invitrogen G21234; RRID:AB_2536530; 1:3000 to 1:5000). Anti-MCU antiserum was raised in rabbits against a KLH-conjugated C-terminal peptide, RTQENTPPTLTEEKAERKY (Pepceuticals, 1:1000). For immunohistochemistry, tissues were incubated with mouse anti-ATP5A (Abcam ab14748; RRID:AB_301447; 1:500) and secondary antibody anti-mouse AF488 (Invitrogen: A11001; RRID:AB_2534069).

### Microscopy

Indirect flight muscle was dissected and fixed in 4% formaldehyde (Agar scientific; R1926) in PBS for 30 minutes, washed twice with PBS, and mounted on slides in Prolong Diamond Antifade mounting medium (Thermo Fisher Scientific; RRID:SCR_015961). Larval epidermal cells were prepared as previously described [33]. Larvae were dissected in PBS and fixed in 4% formaldehyde, for 30 min, permeabilized in 0.3% Triton X-100 for 30 min, and blocked with 0.3% Triton X-100 plus 1% bovine serum albumin in PBS for 1 h at room temperature. Tissues were incubated with ATP5A antibody diluted in 0.3% Triton X-100 plus 1% bovine serum albumin in PBS overnight at 4°C, rinsed three times 10 min with 0.3% Triton X-100 in PBS, and incubated with the appropriate fluorescent secondary antibodies for 2 h at room temperature. The tissues were washed twice in PBS and mounted on slides using Prolong Diamond Antifade mounting medium (Thermo Fisher Scientific). Fluorescence imaging was conducted with a Zeiss LSM 880 confocal microscope/Nikon Plan-Apochromat 63×/1.4 NA oil immersion objective. For adult eyes, images were acquired using a Leica DFC490 camera mounted on a Leica MZ6 stereomicroscope set at maximum zoom.

### Axonal Transport

Analysis of axonal transport was performed on wandering third instar larvae as previously described [34]. Larvae were pinned at each end dorsal side up to a Sylgard (Sigma 761028) slide and cut along the dorsal midline using micro-dissection scissors. Larvae were covered in dissection solution (128 mM NaCl, 1 mM EGTA, 4 mM MgCl_2_, 2 mM KCl, 5 mM HEPES and 36 mM sucrose, adjusted to pH 7 using NaOH), the sides were pinned back and the internal organs removed. Movies were taken using a Nikon E800 microscope with a 60x water immersion lens (NA 1.0 Nikon Fluor WD 2.0) and an LED light source driven by Micromanager 1.4.22 Freeware [35]. A CMOS camera (01-OPTIMOS-F-M-16-C) was used to record 100 frames at a rate of 1 frame per 5 s for *CCAP-GAL4* samples or 1 frame per 2.5 s for *M12-GAL4* samples. Movies were converted into kymographs using Fiji [36] and quantified manually.

### Respirometry Analysis

Respiration was monitored at 30 ^o^C using an Oxygraph-2k high-resolution respirometer (OROBOROS Instruments) using a chamber volume set to 2 mL. Calibration with air-saturated medium was performed daily. Data acquisition and analysis were carried out using Datlab software (OROBOROS Instruments). Five flies per genotype (equal weight) were homogenised in respiration buffer (120 mM sucrose, 50 mM KCl, 20 mM Tris-HCl, 4 mM KH_2_PO_4_, 2 mM MgCl_2_, and 1 mM EGTA, 1 g/l fatty acid-free BSA, pH 7.2). For *MICU3*^27^ experiments, 15 heads per genotype were used. For coupled (state 3) assays, complex I-linked respiration was measured at saturating concentrations of malate (2 mM), glutamate (10 mM) and adenosine diphosphate (ADP, 2.5 mM). Complex II-linked respiration was assayed in respiration buffer supplemented with 0.15 μM rotenone, 10 mM succinate and 2.5 mM ADP.

### ATP Levels

The ATP assay was performed as described previously [37]. Briefly, five male flies of the indicated age or 10 larvae for each genotype were homogenized in 100 μL 6 M guanidine-Tris/EDTA extraction buffer and subjected to rapid freezing in liquid nitrogen. Homogenates were diluted 1/100 with the extraction buffer and mixed with the luminescent solution (CellTiter-Glo Luminescent Cell Viability Assay, Promega). Luminescence was measured with a SpectraMax Gemini XPS luminometer (Molecular Devices). The average luminescent signal from technical triplicates was expressed relative to protein levels, quantified using the DC Protein Assay kit (Bio-Rad). Data from 2-4 independent experiments were averaged and the luminescence expressed as a percentage of the control.

### RNA extraction, cDNA synthesis and real-time PCR

Isolation of total RNA was performed using the RNeasy RNA purification kit (QIAGEN); cDNA was synthesised from total RNA using ‘ProtoScript^®^ II first strand cDNA Synthesis Kit (New England BioLabs, E6560S) according to manufacturer’s instructions. Total RNA concentration was ascertained spectrophotometrically, and equivalent amounts of total RNA underwent reverse transcription for each sample. Quantitative real-time PCR (qRT-PCR) was performed on a CFX96 Touch™ Real-Time PCR Detection System. Gene-specific primers were designed to have oligos spanning an intron whenever possible and their sequence is reported in Supplementary table 1. Carryover DNA was removed with Turbo DNase free (Ambion, Cat. No. AM1907) according to the manufacturer’s protocol. The relative transcript levels of each target gene were normalized against *RpL32* mRNA levels; quantification was performed using the comparative C_T_ method [38].

### Statistical Analysis

Data are reported as mean ± SD, SEM or 95% CI as indicated in figure legends. For climbing analysis, Kruskal-Wallis non-parametric test with Dunn’s post-hoc correction for multiple comparisons was used. For lifespan experiments, significance levels were determined by log-rank tests and reported in the figure legends. Mitochondrial transport was analysed by one-way ANOVA, and ATP and Oroboros measurements analysed by two-tailed t-test. Unless specifically indicated, no significant difference was found between a sample and any other sample in the analysis. Analyses were performed using GraphPad Prism 7 software (RRID:SCR_002798).

## Acknowledgements

This work is supported by MRC core funding (MC_UU_00015/4, MC-A070-5PSB0 and MC_UU_00015/6) and ERC Starting grant (DYNAMITO; 309742). T.P.G. and J.J.L. are supported by an MRC Studentships awarded via the MRC MBU. V.L.H was funded by an EMBO Long-Term Fellowship (ALTF 740-2015) co-funded by the European Commission FP7 (Marie Curie Actions, LTFCOFUND2013, GA-2013-609409). Stocks were obtained from the Bloomington *Drosophila* Stock Center which is supported by grant NIH P40OD018537, and material was obtained from the Drosophila Genomics Resource Center, which is supported by NIH grant 2P40OD010949. We thank Sarah Alam and Ivana Giunta for technical help and all members of the Whitworth lab for discussions.

## Author Contributions

R.T., T.P.G., V.L.H., J.J.L., A.S-M. designed and performed experiments, and analysed the data.

S.v.S. designed, performed and analysed the Ca^2+^ uptake assay with supervision from E.Z.

R.T., T.P.G. and A.J.W. wrote the manuscript with input from all authors.

A.J.W. conceived the study, designed and performed experiments, analysed the data and supervised the work.

